# COMPARATIVE METAGENOMIC ASSESSMENT OF SHORT- AND LONG-READ SEQUENCING TECHNOLOGIES REVEALS UNKNOWN MICROBIAL INFORMATION IN A COMPLEX ENVIRONMENTAL SAMPLE

**DOI:** 10.64898/2025.12.22.696079

**Authors:** Rubén Díaz-Rúa, Daniela I Drautz-Moses, Xiang Zhao, Sadhasivam Perumal, Luke Esau, Angel Angelov, Alexander Putra, Patrick Driguez, Ming Sin Cheung, Emanuele Palescandolo

## Abstract

Metagenomics enables comprehensive exploration of microbial communities but is influenced by library preparation and sequencing technologies, affecting recovery of microbial genomes and proteins. Here, we benchmarked six Illumina short-read library kits at 2×150 bp and 2×250 bp read lengths alongside PacBio HiFi long-read sequencing using a complex environmental sample. Longer short reads (2×250 bp) combined with optimal library preparation approaches notably improved assembly quality, protein detection, and metagenome-assembled genome (MAG) recovery, achieving results similar to those of long-read sequencing. Although long reads yield more contiguous and complete genomes, longer short reads offer a cost-effective, scalable alternative for uncovering microbial and functional diversity. These findings provide critical guidance for metagenomic experimental design, demonstrating the importance of strategic selection of library preparation chemistry and sequencing parameters in revealing unknown microbial information in complex biomes without requiring additional sequencing depth.

**IMPORTANCE:** Metagenomic outcomes are strongly influenced by library preparation and sequencing strategies, yet their combined effects in complex environmental samples remain poorly defined. Here, we provide the first direct comparison of Illumina NovaSeq short-read metagenomic sequencing at 2×150 bp and 2×250 bp across multiple library preparation kits, alongside PacBio long-read sequencing. We show that sequencing read length and library preparation critically shape assembly quality, protein recovery, and metagenome-assembled genome (MAG) reconstruction. Optimized short-read sequencing at 2×250 bp recovered high-quality MAGs approaching those obtained with long-read technologies while substantially improving protein discovery compared to 2×150 bp at the same sequencing depth. Together, these results provide actionable guidance for experimental design and reveal how sequencing read length can be leveraged to improve the recovery of microbial and functional diversity in complex biomes.

## INTRODUCTION

Sequencing technologies have changed the landscape of life sciences research, ushering in a new era of genomic discovery (Koboldt et al. 2013). These advancements have also significantly enhanced our capacity to address ecological and evolutionary questions, enabling comprehensive genetic surveys across the entire tree of life (Burki et al. 2020). The application of high-throughput sequencing for studying microbial communities has given rise to a novel scientific field known as metagenomics, a discipline dedicated to analyzing the collective genomes of organisms coexisting within a given environment. At its core, metagenomics explores genetic material recovered directly from environmental samples, providing a holistic view of entire microbial communities (Handelsman et al. 1998; Alves et al. 2018). Metagenomics offers a snapshot of the true genetic diversity within an ecosystem (Stein et al. 1996) and captures microbial life in its natural habitats, from highly diverse microbial populations in soil (Daniel 2005) to the intricacies of the human gut microbiome (Almeida et al. 2021). This approach bypasses the need to isolate and cultivate individual microorganisms, including the unknown and uncultivated fraction, termed the “dark matter” (Pavlopoulos et al. 2023).

Harvesting microbial information from metagenomes has significantly contributed to the discovery of novel enzymes (Berini et al. 2017), drug candidates from marine products (Trindade et al. 2015), and industrial applications (Lorenz and Eck 2005).

However, variations in experimental protocols are known confounding variables to outcomes of metagenomic studies. Factors such as DNA extraction methods (Costea et al. 2017), library preparation approaches (Jones et al. 2015), sequencing platforms (Chen et al. 2022), sequencing depth and read length can significantly influence the quantitative and qualitative characterization of microbial communities. This underscores the need for standardized and robust protocols to ensure comparability across studies, with library construction method playing a critical role (Jones et al. 2015; Sato et al. 2019). While previous studies testing library preparation methods have largely emphasized differences in sequencing (e.g., GC content, PCR duplicates) and taxonomic traits, identification of novel and unknown proteins and the characterization of non-cultivable microorganisms are often not adequately addressed.

Sequencing technologies are commonly classified as (i) short-read and high-throughput (<300 bp) or Next Generation Sequencing (NGS), and (ii) long-read (> 300 bp) with the later facing limitations in throughput, cost, and DNA quality and quantity requirements (Kim et al. 2024), but often lead to improved genome assembly (Gehrig et al. 2022). Although 2×150 bp reads are widely employed in metagenomics, and prior studies have examined the impact of read length through simulations and computational comparisons (Kang et al. 2022; Treiber et al. 2020), empirical, platform-specific assessments of metagenomics performance comparing different read lengths by sequencing the same library under otherwise identical conditions on the Illumina NovaSeq 6000 remain scarce.

Furthermore, variations in environmental sample composition and experimental protocols pose a significant challenge to reproducibility. In comparative studies, while commercial mock bacterial communities are useful for benchmarking purposes, their limited species diversity compared to natural microbiomes may not be sufficient to detect protocol differences (Gaulke et al. 2021). In addition, the matrix in which microbial communities are often embedded in can add additional challenges to successful sample processing and reproducibility. Sanders et al. emphasized that authentic environmental samples provide a more realistic framework for evaluating library preparation methods and sequencing technologies. They used long-read sequencing as a benchmarking in environments permitting a realistic assessment of sequencing and assembly strategies for novel and complex microbial communities (Sanders et al. 2019).

In this study, we conducted comparative benchmarking of metagenomic sequencing strategies to assess the impact of library preparation, read length and sequencing technology on the recovery of microbial information from complex environmental communities. We systematically compared six Illumina short-read (SRS) library preparation kits, each tested at two sequencing read lengths (2×150 bp and 2×250 bp), along with a PacBio HiFi long-read (LRS) dataset, covering a total of thirteen distinct sequencing conditions. This design enabled us to assess the impact of library construction, sequencing read length, and technology on taxonomic profiling, protein detection, and the recovery of high-quality metagenome-assembled genomes (MAGs). By integrating these results, we aimed to identify sequencing strategies that maximize the recovery of taxonomic, functional, and genome-resolved microbial information from complex metagenomes.

## MATERIALS AND METHODS

### Sample Preparation and DNA Extraction

A unique and highly diverse mock environmental sample was engineered to provide sufficient DNA quantity and quality for comprehensive testing across various library preparation kits and sequencing technologies without being influenced by different sampling sites and DNA extraction. This approach also allowed for comprehensive testing across various conditions and platforms, using a specimen similar to actual environmental samples rather than relying on mock and commercial material (Sanders et al. 2019).

For this, samples were collected from two distinct environments: marine mangrove sediments and terrestrial palm tree soil, both expected to be rich in microbial diversity. For each environment, five DNA extractions were carried out with the DNeasy PowerMax Soil Kit (Qiagen, Hilden, Germany), following the manufacturer’s instructions and using 8 grams of soil as input material for each extraction. The large amount of material used per replicate ensured a high representation of microbes from both sites. Subsequently, the DNA aliquots from all extractions were pooled equimolarly in a single aliquot, which was then cleaned and concentrated using the NucleoSpin gDNA Clean-up Kit (MACHEREY-NAGEL, Dueren, Germany).

### Library Preparation and Sequencing

For short-read sequencing (SRS), the following library preparation kits were used: NEBNext Ultra II DNA Library Prep Kit for Illumina (NEB, Ipswich, MA, USA), Nextera XT DNA Library Preparation Kit (Illumina, San Diego, CA, USA), Ovation Ultralow System V2 (Tecan, Männedorf, Switzerland) and TruSeq DNA Nano (Illumina, San Diego, CA, USA). Libraries were prepared in triplicate following the manufacturers’ instructions, and when the protocol allowed, we tested different DNA inputs, 1 and 100 ng (see Table 1 for all library preparation details). Size distribution was assessed using the Bioanalyzer 2100 (Agilent Technologies, Santa Clara, CA, USA). The libraries were then quantified using the KAPA Library Quantification Kit for Illumina Platforms (Roche, Basel, Switzerland). After equimolar pooling of all libraries, 1 nM aliquots were sequenced on NovaSeq6000 SP flow cells at read lengths of 2×150 bp and 2×250 bp (referred hereinafter as 150PE and 250PE, respectively).

**Table 1.** Library preparation kits used in this study.

| Commercial kit name | Sample name (replicate) | Co-assembly name | Fragmentation method | Input (ng) | Estimated price per library (\$) | Assay time (hours) |
| --- | --- | --- | --- | --- | --- | --- |
| NEBNext® Ultra™ II DNA Library Prep Kit for Illumina® | NEB-1_1 | NEB-1 | Enzymatic | 1 | 40 | 6 |
|  | NEB-1_2 |  |  |  |  |  |
|  | NEB-1_3 |  |  |  |  |  |
| NEBNext® Ultra™ II DNA Library Prep Kit for Illumina® | NEB-100_1 | NEB-100 | Enzymatic | 100 | 40 | 6 |
|  | NEB-100_2 |  |  |  |  |  |
|  | NEB-100_3 |  |  |  |  |  |
| Nextera® XT DNA Library Preparation Kit | NEXT-1_1 | NEXT-1 | Transposome | 1 | 65 | 5 |
|  | NEXT-1_2 |  |  |  |  |  |
|  | NEXT-1_3 |  |  |  |  |  |
| Ovation® Ultralow System V2 | Ovation-1_1 | Ovation-1 | Sonication | 1 | 54 | 6 |
|  | Ovation-1_2 |  |  |  |  |  |
|  | Ovation-1_3 |  |  |  |  |  |
| Ovation® Ultralow System V2 | Ovation-100_1 | Ovation-100 | Sonication | 100 | 54 | 6 |
|  | Ovation-100_2 |  |  |  |  |  |
|  | Ovation-100_3 |  |  |  |  |  |
| TruSeq® DNA Nano | TruSeq-100_1 | TruSeq-100 | Sonication | 100 | 50 | 7 |
|  | TruSeq-100_2 |  |  |  |  |  |
|  | TruSeq-100_3 |  |  |  |  |  |
| Metagenomics Shotgun Library Preparation Using SMRTbell Express Template Prep Kit 2.0 | PacBio | PacBio | Shearing | 2,000 | 552 | 10 |

For long-read sequencing (LRS), 0.45x Ampure PB Beads (PacBio, Menlo Park, CA, USA) were used to concentrate the gDNA. DNA concentration and size distribution were determined using the Qubit HS dsDNA Assay Kit with the Qubit 2.0 Fluorometer (Life Technologies, CA, USA) and the Femto Pulse platform (Agilent Technologies, Santa Clara, CA, USA), respectively. The library was prepared with 2,000 ng of input DNA using the SMRTbell Express Template Prep Kit 2.0 (PacBio, Menlo Park, CA, USA). Final library concentration was measured using the Qubit dsDNA High Sensitivity Kit and library size distribution was assessed with the Agilent Femto Pulse. Next, 100 pM of the SMRTbell library was mixed with the sequencing primer and Sequel II DNA polymerase from the Sequel II Binding kit 2.2 (101-894-200), following the SMRTlink sample set-up guidelines. HiFi sequencing was then performed with a 30-hour movie time in adaptive loading mode, using the Sequel II Sequencing Kit 2.0 (101-820-200) and SMRT Cell 8M Tray (101-389-001).

### Sequencing Data Analysis

Sequencing data were processed using standard metagenomic pipelines with specific tools and packages for quality control, adapter trimming, genome assembly, gene prediction, protein annotation, binning, and MAG characterization. Figure 1 provides an overview of the short- and long-read analyses pipelines.

**Figure 1.**
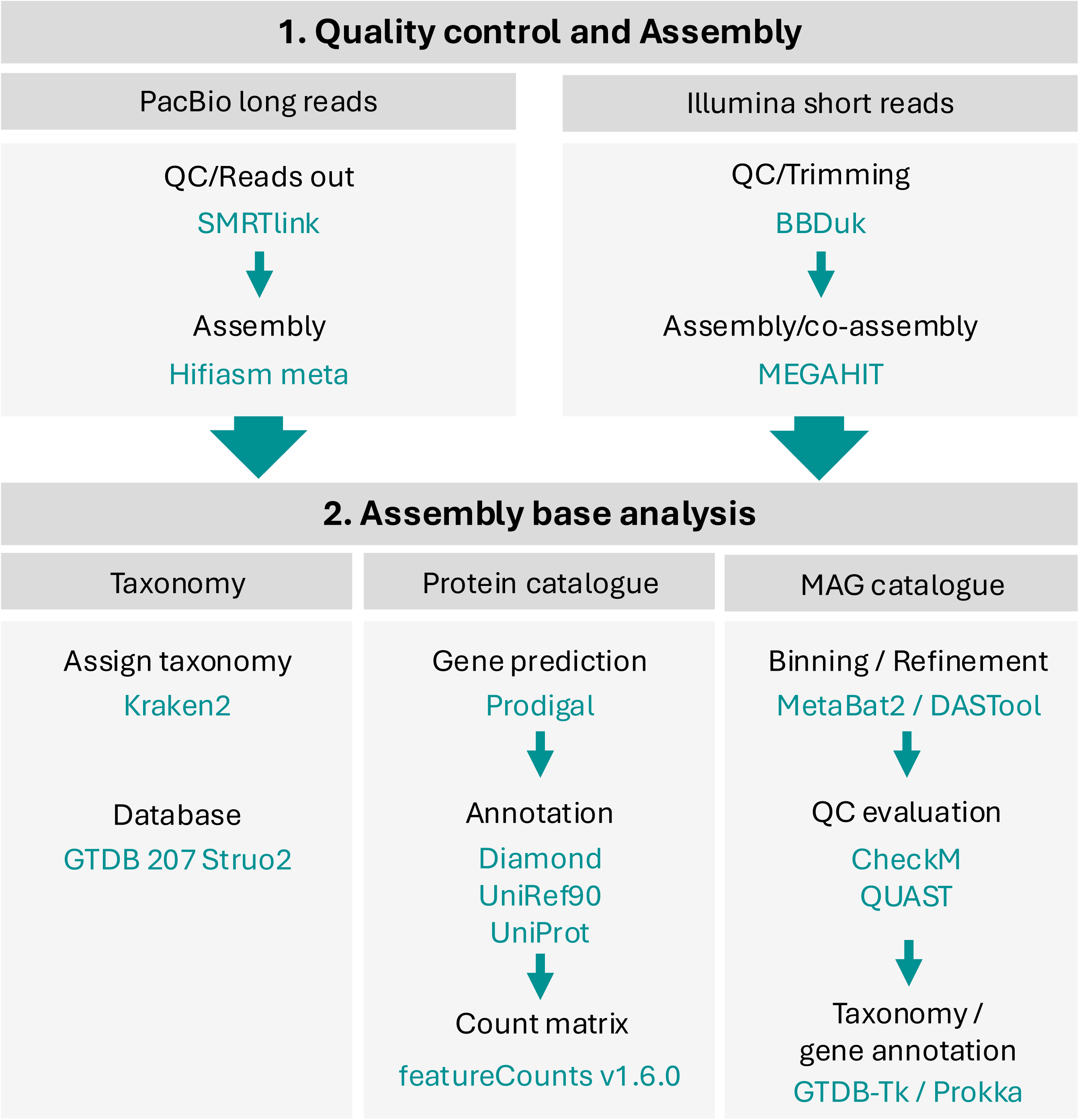
Overview of the sequencing data analysis pipeline used for both sequencing technologies.

### Illumina Short-Read Quality Check, Trimming and Assembly

BBDuk from the BBTools package was used to trim adapters and remove low-quality sequences. Random down-sampling was performed to ensure an equal number of reads across samples was used for subsequent analyses. Down-sampled data were pooled per replicate, referred to as ‘kit merge’ (KM), or pooled across all SRS kits and termed ‘global merge’ (GM). High-quality reads were assembled using MEGAHIT (Li et al. 2015) with the option “--presets meta-large”. Assembly statistics, such as number of contigs, contig lengths, and N50/N90 metrics, were calculated using assembly-stats (GitHub repository sanger-pathogens/assembly-stats). Reads were mapped to the contigs using BWA (Li and Durbin 2009).

### PacBio Long-Read Quality Check, Trimming and Assembly

PacBio HiFi reads were quality-checked and filtered using the SMRTlink software (PacBio, Menlo Park, CA, USA). Assembly was performed using the hifiasm meta-assembler (Feng et al. 2022) through the official PacBio pipeline (GitHub repository PacificBiosciences/pb-metagenomics-tools).

### Taxonomic Assignment and Diversity Analyses

Taxonomic classification for Illumina and PacBio contigs was performed using the Kraken algorithm (Wood et al. 2019) within OmicsBox V3.4 (BioBam Bioinformatics, 2019) and the GTDB 207 Struo2 database (Parks et al. 2022). Beta diversity was assessed through Principal Coordinate Analysis (PCoA) using Bray-Curtis dissimilarity matrix on the OTU matrix at genus level for the SRS replicates. Significance of the resulting ordination was tested with Permutational MANOVA (PerMANOVA) and 999 permutations (adonis2, vegan package in R). Genus-level richness was calculated using the *estimate_richness* function from the *phyloseq* package (McMurdie and Holmes 2013) in R 4.4.0 (“RCoreTeam, 2024,” n.d.).

### Differential Abundance Analysis of Taxa

Differential abundance analysis at genus level was carried out with the edgeR package (Robinson et al. 2010) in OmicsBox V3.4, applying a Binomial Generalized Linear Model (Binomial GLM) with TMMwsp normalization using a minimum of 10 counts per million (CPM) and a minimum of 5 counts per sample to exclude low-abundance taxa that could overestimate the differences. Statistical differences between the groups were assessed using the Student’s t-test with multiple testing correction (False Discovery Rate, FDR) as previously described, (Benjamini and Hochberg 1995). Corrected P-values of <0.05 and with a log-fold change of ±1 were considered statistically significant. Significant genera were visualized in a heatmap where CPM values were presented on a logarithmic scale as Z-scores (Skuta et al. 2014).

### Gene Prediction, Annotation, and Protein Catalogue

For both SRS and LRS data sets, gene prediction was conducted with Prodigal (Hyatt et al. 2010) with the option “-p meta”, followed by a dereplication step of clustering at 95% amino acid identity with CD-HIT v4.8.1 (Fu et al. 2012) to remove genetic redundancy.

Subsequently, protein sequences obtained by CD-HIT were annotated against the UniRef90 database using Diamond blastp (Buchfink et al. 2015), requiring a minimum read coverage of 90% and a minimum identity of 80%.

In order to decrease fragmented proteins that could inflate the results in large datasets as the KM and GM co-assemblies, a more stringent filtering process was performed. Only contigs larger than 500 bp and genes longer than 100 bp (33 amino acids) were selectively retrieved. A unique set of proteins was identified using *featureCounts* v1.6.0 (Liao et al. 2014) to count the number of reads that mapped to each predicted protein. Proteins with a hit against the UniRef90 database (identity 80%) were then extracted, aggregating counts for proteins with the same UniProt IDs and keeping only proteins with more than one read. Protein hit information was downloaded from UniProt using UPIMAPI (Sequeira et al. 2022). To assess the number of unique annotated and uncharacterized proteins, and to focus specifically on those with identified functions, a non-redundant protein catalogue was generated by collapsing protein counts across the KM and GM datasets based on shared protein identifiers. Given its substantially higher sequencing depth—approximately six times greater than that of the KM co-assemblies—and its inclusion of reads from all Illumina library preparation kits, the GM dataset was selected as the reference protein database for this study.

### Principal Coordinate Analysis (PCoA) for Identified Proteins in the Illumina Kits

PCoA was performed on normalized gene count matrices (Bray-Curtis distances) using featureCounts v1.6.0 (Liao et al. 2014), with the GTF and BAM files for the individual samples obtained with Prodigal. The individual BAM files were generated by mapping the raw reads against the GM co-assembly. The count matrix was then normalized using the Variance Stabilizing Transformation (VST). Genes with 0 variances were filtered out. Principal Components were generated using the *prcomp* function in R on the Bray-Curtis distance. Statistical significance was tested with PerMANOVA.

### MAG Reconstruction, Classification, and Comparison

For MAG reconstruction, a binning process was carried out for the KM and GM co-assemblies and the PacBio dataset. MetaBAT2 (Kang et al. 2019) was used for this purpose, with a minimum contig length of 1,500 bp. After the binning, a refinement step was applied with DAS Tool (Sieber et al. 2018). Quality checks were performed with CheckM (Parks et al. 2015). PacBio bins were used as internal reference genomes, further serving as a benchmark for various short-read sequencing strategies. Taxonomic classification of high-quality MAGs (completeness > 90%, contamination <5%) (Bowers et al. 2017) was obtained with the Genome Taxonomy Database Toolkit (GTDB-Tk) (Chaumeil et al. 2019). For high-quality PacBio MAGs with Average Nucleotide Identities (ANIs) <95% (standard speciation threshold) (Jain et al. 2018), a phylogenetic tree was constructed with the GTDB-Tk *de novo* workflow (de_novo_wf) in order to detect potential new microbes. The outgroup taxon (--outgroup_taxon) option was used to reduce the tree size (Supplementary Figure 3). For MAG comparison between sequencing technologies, the ANI between the SRS and LRS bins was determined using FastANI v1.32 (Jain et al. 2018), and the MASH distance and p-value were calculated to assess MAG similarity (Ondov et al. 2016). QUAST was used to determine assembly metrics (Gurevich et al. 2013), and Prokka for gene annotation (Seemann 2014). Visualization of each *de novo* MAG assembly was conducted with Bandage (Wick et al. 2015). Orthologous gene clusters among the selected bins were identified using OrthoVenn3 (Sun et al. 2023). Protein sequences (in FASTA format) from each bin were uploaded to the web server with default parameters. The analysis produced clusters of shared and unique orthologous. Finally, to explore the potential novel function of these high-quality MAGs, biosynthetic gene clusters (BGCs) were identified using antiSMASH version 8.0 (Blin et al. 2025) with default parameters. The tool was run in *“full mode”* to detect known and putative secondary metabolite clusters, including those for polyketides (PKS), nonribosomal peptides (NRPS), terpenes, and others. Cluster boundaries, annotations, and predicted products were generated based on comparison with the MIBiG reference database (Zdouc et al. 2025).

### Statistical Analysis and Graphing

General statistical analysis and bar plots were performed using GraphPad Prism 9.0 for Mac. ANOVA followed by Tukey’s test was used to compare SRS library preparation kits. Student’s t-test was applied to compare the 150PE and 250PE sequencing read lengths. All differences were considered significant when p < 0.05.

### Data availability

All sequencing reads were deposited in the ENA database under project accession numbers PRJEB50284.

## RESULTS

### DNA Isolation, Illumina Library Preparation and Sequencing Results

DNA extraction from the engineered terrestrial soil and marine sediment mixture yielded a total of 6.2 µg DNA. Yield and DNA fragment size were sufficient for evaluating multiple SRS library preparation kits, sequencing lengths, and LRS.

Library yield and size distribution were assessed to compare SRS kit performance. Each SRS kit showed characteristic library yields and size distributions (Supplementary Figure 1). At 1 ng DNA input, the Ovation kit produced the highest library yield, followed by Nextera and NEB. At 100 ng DNA input, TruSeq resulted in a narrower library fragment distribution, followed by Ovation and NEB. Following manufacturer’s recommendation, for 1 ng of DNA input, the NEB kit required fewer PCR cycles than Ovation and Nextera. This could explain the higher PCR duplication rates and higher library yields in the Ovation and Nextera data sets. Of note, the fewest PCR duplicates were generated with the TruSeq kit (Table 2), even though this kit did not require the lowest PCR cycles (Supplementary Figure 1).

**Table 2.**
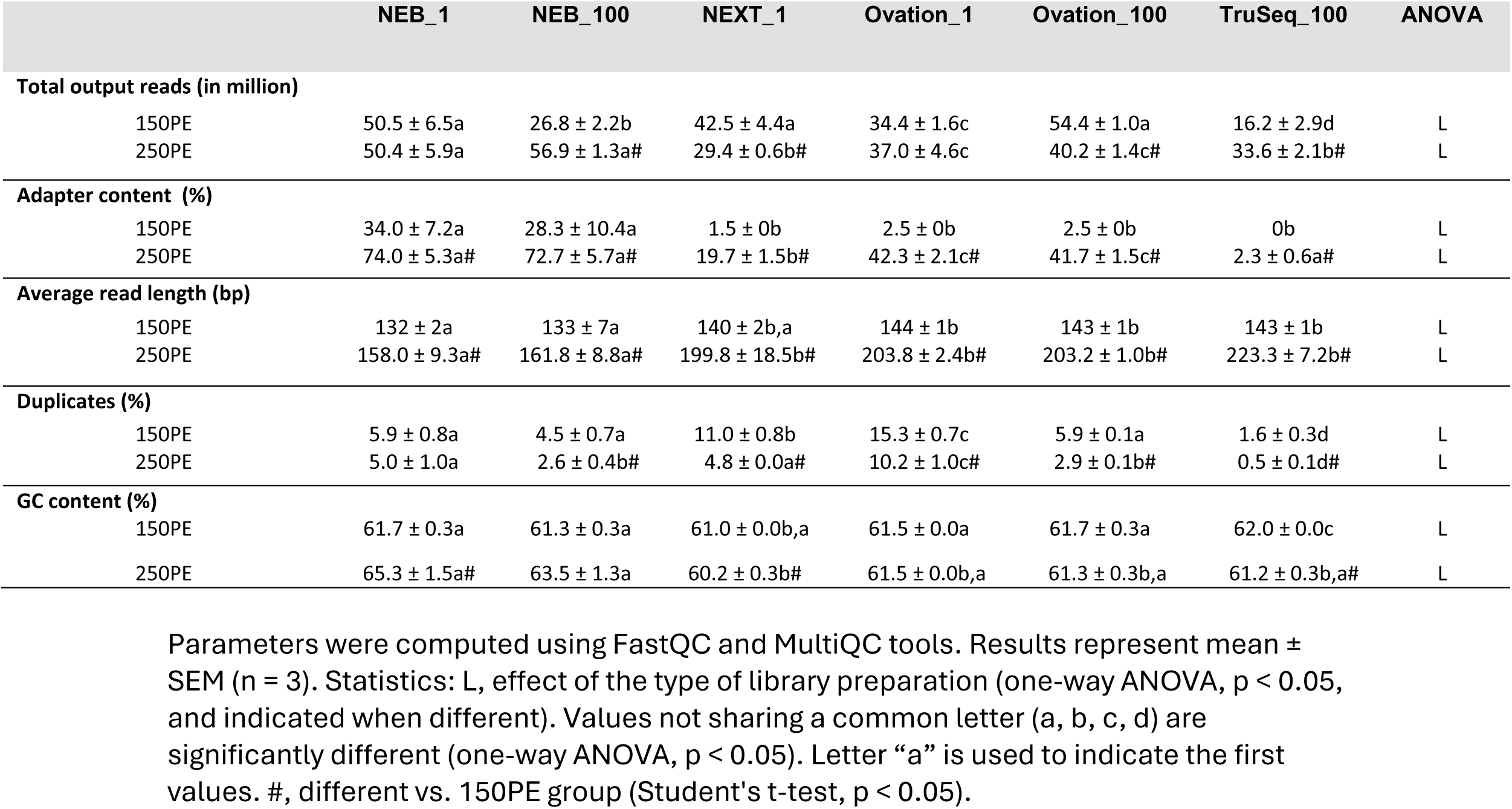
SRS Illumina sequencing results per replicate. Total output reads adapter content, average read length, duplicate sequences and GC content of the different Illumina library preparation kits sequenced by 150PE or 250PE read length mode.

We observed that shorter fragments (Ovation and NEB libraries), clustered more efficiently than the longer fragments of the Nextera and TruSeq libraries (Table 2). The latter produced three times less data in the 150PE run than the highest-yielding library despite equimolar loading concentrations. Nextera libraries displayed a size distribution similar to Ovation but with enrichment in longer fragments (Supplementary Figure 1). Expectedly, libraries with shorter fragment sizes showed higher adapter content, particularly in the 250PE run, where longer read lengths increased the likelihood of sequencing into the adapters (Table 2).

After adapter trimming and quality filtering, the 150PE dataset indicated that NEB libraries had an average effective insert length ∼10 bp shorter than those prepared with the other kits, for both the 1 ng and 100 ng DNA inputs. For the 250PE run, TruSeq achieved the longest average effective insert length (223 bp), followed by Ovation (203 bp for both DNA inputs), Nextera (199 bp), and NEB (158-162 bp) (Table 2).

### Impact of Sequencing Read Length on Assembly Metrics

For the SRS datasets, after trimming and subsampling, *de novo* assemblies were generated for each replicate and assembly metrics were compared (Table 3). Overall, NEB libraries showed the lowest N50, N90, and average contig lengths, while TruSeq and Nextera showed the highest values. The 250PE run showed significant improvements in all assembly metrics, including contig counts and gene recovery when compared to the 150PE sequencing mode (Tables 3 and 4), particularly for TruSeq and Nextera libraries, which showed the larger effective insert sizes.

**Table 3.** SRS Illumina assembly results per replicate. Summatory bases of the assembly, number of contigs, average of contigs, largest contigs, N50 and N50 values of the different Illumina library preparation kits sequenced by 150PE or 250PE read length mode.

|  | NEB_1 | NEB_100 | NEXT_1 | Ovation_1 | Ovation_100 | TruSeq_100 | ANOVA |
| --- | --- | --- | --- | --- | --- | --- | --- |
| **Sum of assembly (Mbp)** |  |  |  |  |  |  |  |
| 150PE | 93 ± 2a | 111 ± 13b | 141 ± 3c | 139 ± 2c | 144 ± 5c | 129 ± 4b,c | L |
| 250PE | 438.5 ± 3.5a# | 472 ± 71a# | 1637 ± 52b# | 1057 ± 6c# | 1100 ± 50c# | 2605 ± 34d# | L |
| **Contigs (10<sup>6</sup>)** |  |  |  |  |  |  |  |
| 150PE | 2.4 ± 0.05a | 2.8 ± 0.3b | 3.3 ± 0.1c | 3.4 ± 0.05c | 3.5 ± 0.1c | 2.9 ± 0.1c,b | L |
| 250PE | 11.6 ± 0.1a# | 12.5 ± 1.8a# | 40.7 ± 1.1b# | 27.0 ± 0.2c# | 28.0 ± 1.2c# | 62.6 ± 1.3d# | L |
| **Average (bp)** |  |  |  |  |  |  |  |
| 150PE | 392 ± 0.2a | 394 ± 3.3a | 427 ± 1.1b | 409 ± 0.1c | 410 ± 0.8c | 451 ± 1.4d | L |
| 250PE | 377.2 ± 0.3a# | 376 ± 1.5a# | 402 ± 1.7b# | 392 ± 0.1c# | 393 ± 0.5c# | 416 ± 3.6d# | L |
| **Largest (Mbp)** |  |  |  |  |  |  |  |
| 150PE | 14 ± 0.5a | 25 ± 0.4b | 30 ± 2.6b | 31 ± 3.5b | 32 ± 5.3b | 29 ± 6.4b | L |
| 250PE | 26 ± 0.4# | 25 ± 4.2 | 27 ± 2.2 | 25 ± 0.8 | 29 ± 2.9 | 25 ± 0.8 |  |
| **N50** |  |  |  |  |  |  |  |
| 150PE | 373 ± 0a | 375 ± 3a | 402 ± 1b | 386 ± 0c | 386 ± 1c | 425 ± 1d | L |
| 250PE | 363.0 ± 2.6a# | 360 ± 2a# | 388 ± 2b# | 380 ± 1c# | 381 ± 1c# | 402 ± 4d# | L |
| **N90** |  |  |  |  |  |  |  |
| 150PE | 312 ± 0a | 313 ± 1a | 316 ± 0b | 314 ± 0c | 314 ± 0c | 317 ± 1d | L |
| 250PE | 310.3 ± 0.6a# | 310 ± 0a# | 318 ± 1b# | 314 ± 0c | 314 ± 0c | 329 ± 2d# | L |

Parameters were calculated with the software assembly-stats (GitHub repository sanger-pathogens / assembly-stats). Results represent mean ± SEM (n = 3). Statistics: L, effect of the type of library preparation (one-way ANOVA, p < 0.05, and indicated when different). Values not sharing a common letter (a, b, c, d) are significantly different (one-way ANOVA, p < 0.05). Letter “a” is used to indicate the first values. No letters = no statistical difference. #, different vs. 150PE group (Student's t test, p < 0.05).

**Table 4.**
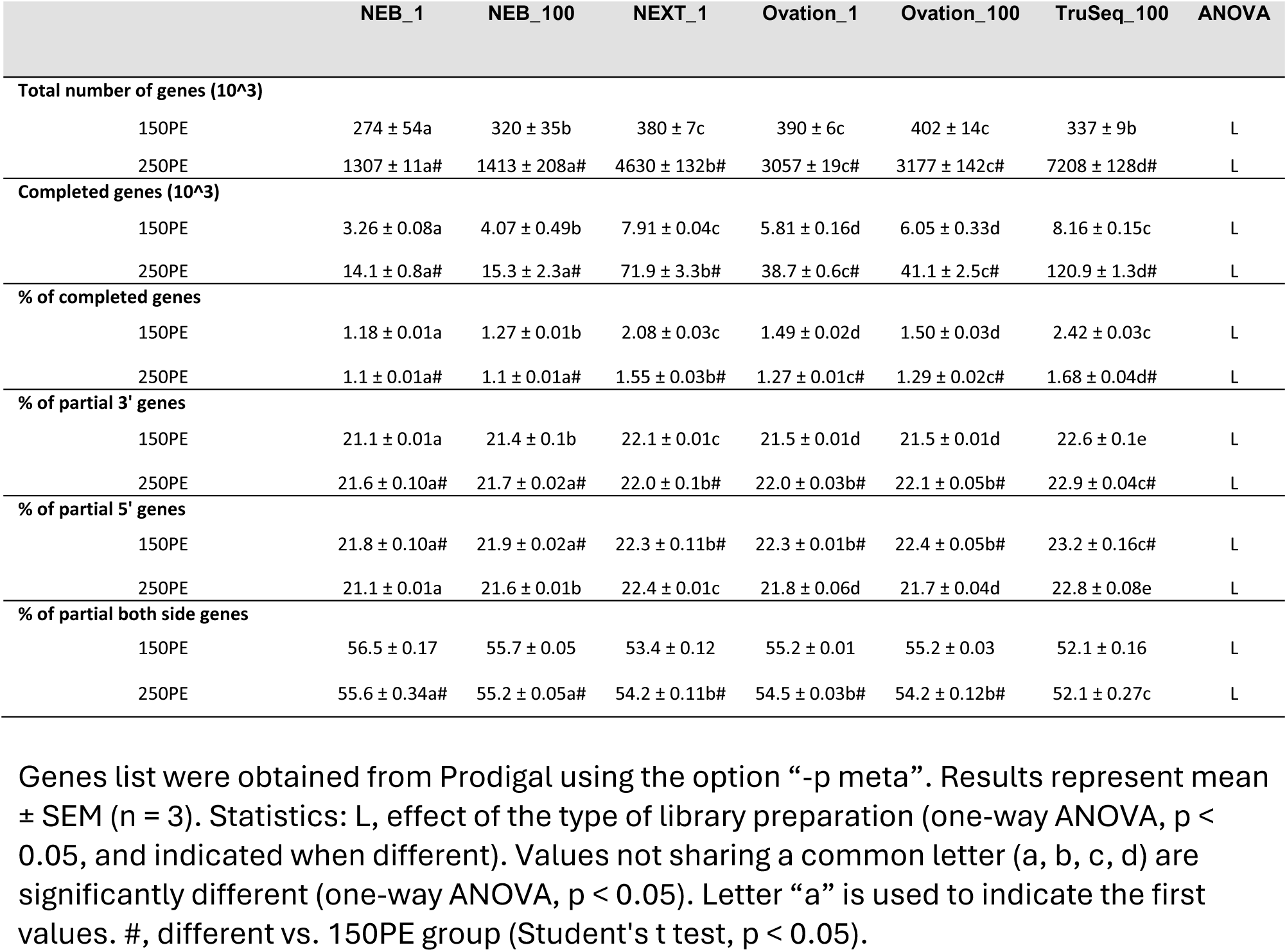
SRS Illumina genes results per replicate. Total number of genes, completed genes, percentage of completed genes, percentage of partial 3’ genes, percentage of partial 5’ genes and percentage of partial both side genes of the different Illumina library preparation kits sequenced by 150PE or 250PE read length mode.

|  | NEB_1 | NEB_100 | NEXT_1 | Ovation_1 | Ovation_100 | TruSeq_100 | ANOVA |
| --- | --- | --- | --- | --- | --- | --- | --- |
| **Total number of genes (10<sup>3</sup>)** |  |  |  |  |  |  |  |
| 150PE | 274 ± 54a | 320 ± 35b | 380 ± 7c | 390 ± 6c | 402 ± 14c | 337 ± 9b | L |
| 250PE | 1307 ± 11a# | 1413 ± 208a# | 4630 ± 132b# | 3057 ± 19c# | 3177 ± 142c# | 7208 ± 128d# | L |
| **Completed genes (10<sup>3</sup>)** |  |  |  |  |  |  |  |
| 150PE | 3.26 ± 0.08a | 4.07 ± 0.49b | 7.91 ± 0.04c | 5.81 ± 0.16d | 6.05 ± 0.33d | 8.16 ± 0.15c | L |
| 250PE | 14.1 ± 0.8a# | 15.3 ± 2.3a# | 71.9 ± 3.3b# | 38.7 ± 0.6c# | 41.1 ± 2.5c# | 120.9 ± 1.3d# | L |
| **% of completed genes** |  |  |  |  |  |  |  |
| 150PE | 1.18 ± 0.01a | 1.27 ± 0.01b | 2.08 ± 0.03c | 1.49 ± 0.02d | 1.50 ± 0.03d | 2.42 ± 0.03c | L |
| 250PE | 1.1 ± 0.01a# | 1.1 ± 0.01a# | 1.55 ± 0.03b# | 1.27 ± 0.01c# | 1.29 ± 0.02c# | 1.68 ± 0.04d# | L |
| **% of partial 3' genes** |  |  |  |  |  |  |  |
| 150PE | 21.1 ± 0.01a | 21.4 ± 0.1b | 22.1 ± 0.01c | 21.5 ± 0.01d | 21.5 ± 0.01d | 22.6 ± 0.1e | L |
| 250PE | 21.6 ± 0.10a# | 21.7 ± 0.02a# | 22.0 ± 0.1b# | 22.0 ± 0.03b# | 22.1 ± 0.05b# | 22.9 ± 0.04c# | L |
| **% of partial 5' genes** |  |  |  |  |  |  |  |
| 150PE | 21.8 ± 0.10a# | 21.9 ± 0.02a# | 22.3 ± 0.11b# | 22.3 ± 0.01b# | 22.4 ± 0.05b# | 23.2 ± 0.16c# | L |
| 250PE | 21.1 ± 0.01a | 21.6 ± 0.01b | 22.4 ± 0.01c | 21.8 ± 0.06d | 21.7 ± 0.04d | 22.8 ± 0.08e | L |
| **% of partial both side genes** |  |  |  |  |  |  |  |
| 150PE | 56.5 ± 0.17 | 55.7 ± 0.05 | 53.4 ± 0.12 | 55.2 ± 0.01 | 55.2 ± 0.03 | 52.1 ± 0.16 | L |
| 250PE | 55.6 ± 0.34a# | 55.2 ± 0.05a# | 54.2 ± 0.11b# | 54.5 ± 0.03b# | 54.2 ± 0.12b# | 52.1 ± 0.27c | L |

Genes list were obtained from Prodigal using the option “-p meta”. Results represent mean ± SEM (n = 3). Statistics: L, effect of the type of library preparation (one-way ANOVA, p < 0.05, and indicated when different). Values not sharing a common letter (a, b, c, d) are significantly different (one-way ANOVA, p < 0.05). Letter “a” is used to indicate the first values. #, different vs. 150PE group (Student's t test, p < 0.05).

Assembly metrics revealed that the GM co-assembly produced more contigs and larger assembly sizes than any of the single KM assemblies (Table 5), due to the higher sequencing depth. The PacBio long reads further improved assembly stats, in particular N50, N90 and contig sizes (Table 5 and Supplementary Figure 2), as expected.

**Table 5.** SRS co-assemblies and LRS assembly results. Number of total reads used, summatory bases of the assembly, number of contigs, average of contigs, largest contigs, N50 and N900 values of the different SRS Illumina library preparation kits sequenced by 150PE or 250PE read length mode.

|  | Number of reads<br>(in million) | Sum of bases<br>(in million) | Number of<br>contigs (10<sup>3</sup>) | Average (bp) | Largest (Mbp) | N50 | N90 |
| --- | --- | --- | --- | --- | --- | --- | --- |
| **PacBio** | 1.7 | 531 | 15 | 35,894 | 8269 | 36,899 | 22,206 |
| **All reads downsampling (GM)** |  |  |  |  |  |  |  |
| 150PE | 216 | 5,288 | 8,426 | 628 | 457 | 642 | 339 |
| 250PE | 216 | 20,448 | 40,048 | 511 | 303 | 465 | 331 |
| **Per kit (KM)** |  |  |  |  |  |  |  |
| **NEB-1** |  |  |  |  |  |  |  |
| 150PE | 38 | 518 | 1,188 | 436 | 26 | 405 | 316 |
| 250PE | 38 | 1,295 | 3,215 | 403 | 26 | 382 | 313 |
| **NEB-100** |  |  |  |  |  |  |  |
| 150PE | 38 | 532 | 1,226 | 434 | 25 | 403 | 315 |
| 250PE | 38 | 1,437 | 3,563 | 403 | 25 | 380 | 312 |
| **NEXT-1** |  |  |  |  |  |  |  |
| 150PE | 38 | 747 | 1,552 | 481 | 29 | 446 | 320 |
| 250PE | 38 | 4,794 | 1,1052 | 434 | 60 | 407 | 321 |
| **Ovation-1** |  |  |  |  |  |  |  |
| 150PE | 38 | 763 | 1,668 | 458 | 49 | 425 | 318 |
| 250PE | 38 | 3,237 | 7,683 | 421 | 49 | 398 | 317 |
| **Ovation-100** |  |  |  |  |  |  |  |
| 150PE | 38 | 787 | 1,717 | 458 | 41 | 426 | 318 |
| 250PE | 38 | 3,348 | 7,935 | 422 | 46 | 399 | 317 |
| **TruSeq-100** |  |  |  |  |  |  |  |
| 150PE | 38 | 731 | 1435 | 510 | 73 | 482 | 322 |
| 250PE | 38 | 7298 | 16253 | 449 | 92 | 418 | 332 |

Parameters were calculated with the software assembly-stats (GitHub repository sanger-pathogens / assembly-stats). Results represent unique value per kit and sequencing length.

### Library Preparation and Sequencing Technologies Affect Taxonomy and Diversity

Principal coordinate analysis (PCoA) of the OTUs at genus-level revealed a significant difference between library preparation kits for both SRS runs, 150PE (adonis2 test, R² = 0.617, p = 0.001) and 250PE (adonis2 test, R² = 0.738, p = 0.001). Nextera libraries formed the most distinct clusters, indicating the largest differences in beta diversity associated with this kit (Figure 2-A1). However, these trends were not statistically significant when tested across all kits using pairwise comparisons (data not shown). To further explore these patterns in ordination, a differential abundance analysis was performed comparing the 150PE and 250PE libraries. We identified 16 (11 over-represented and 5 under-represented) and 12 (2 over-represented and 10 under-represented) genera in Nextera libraries, respectively, when compared to other kits (FDR < 0.05 and a logFC ±1), with the top ten affected taxa shown in Figure 2-A2.

**Figure 2.**
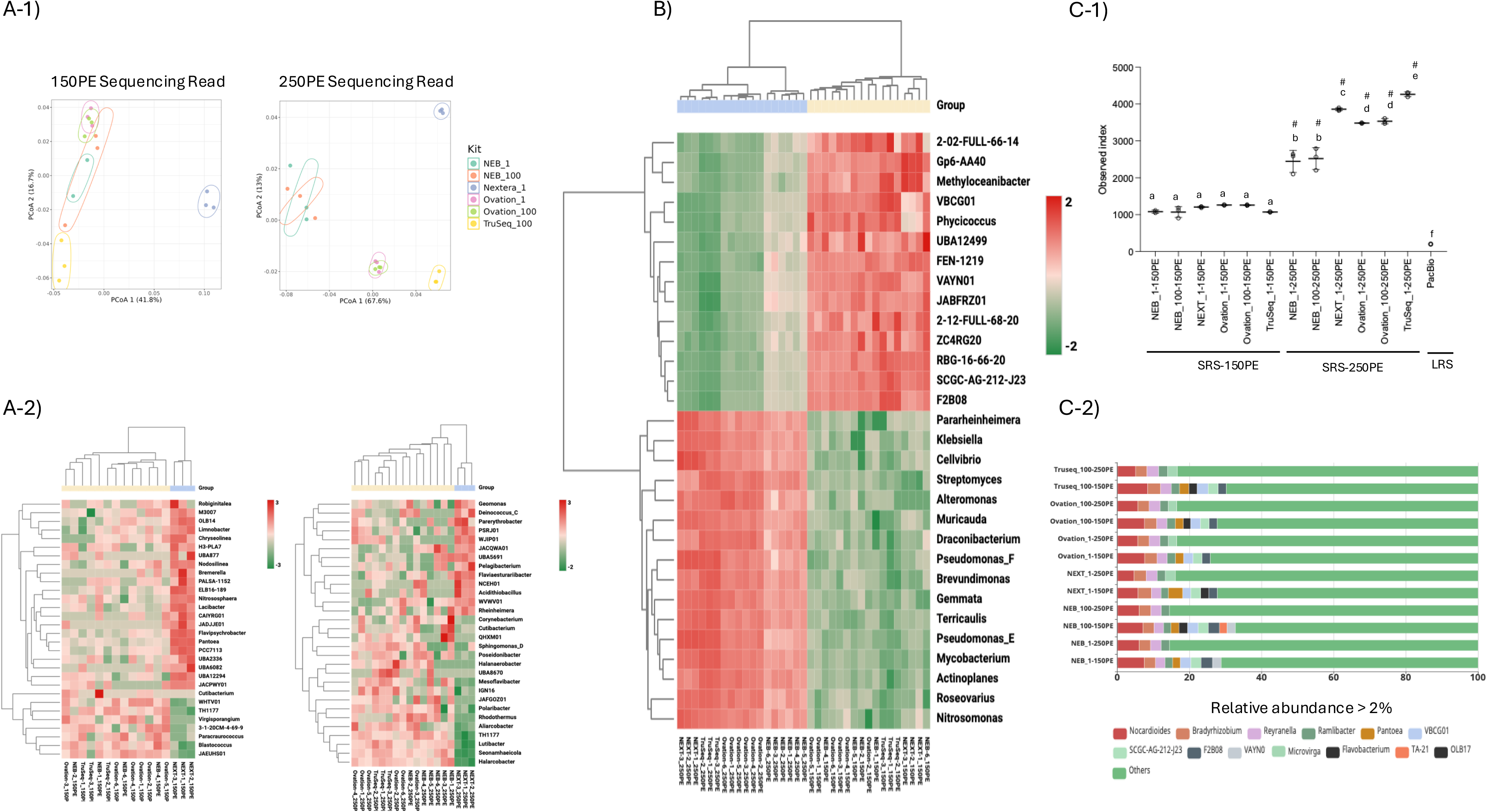
Comparison of library preparation kits and sequencing technologies for taxonomic assignment. A-1) Principal Coordinate Analysis (PCoA) based on Bray-Curtis dissimilarity of detected OTUs for each Illumina library preparation kit and DNA input replicate. Each point represents a single sample and is colored by library preparation kit. Differences in community composition among groups were tested by PERMANOVA (R^2^ and p-value; 999 permutations). A-2) Heatmap of the ten most affected genera in Nextera libraries compared with the other kits, shown for both read lengths. Abundance is displayed as a color gradient. Hierarchical clustering is shown by dendrograms on the left (OTUs) and top (samples) using Euclidean distance. Top annotation bars indicate experimental conditions; OTU names are shown on the right. B) Heatmap of the top 30 affected taxa between the two read lengths, displayed as in A-2). C-1) Alpha diversity (observed richness) at the genus level for each Illumina SRS kit and the PacBio LRS sample. Values are mean ± SEM (n = 3). Statistics: letters indicate differences among library preparation kits (one-way ANOVA, p < 0.05); groups not sharing a letter differ significantly; # indicates differences between 150PE and 250PE (Student’s t-test, p < 0.05). C-2) Stacked bar plot of genus-level relative abundance community composition across Illumina kits and sequencing technologies. Percentages of total classified contigs are shown in parentheses. Genera with <2% relative abundance are grouped as “Others”.

Next, a general differential abundance analysis comparison at the genus level of the two sequencing lengths (150PE vs 250PE) revealed that out of 5,301 genera meeting the criteria of more than 10 CPM, 775 genera were significantly affected (FDR < 0.05 and logFC ±1), with 633 over-represented and 142 under-represented in the 250PE data set. Hierarchical clustering of the top 30 affected taxa showed a clear separation between sequencing lengths, highlighting consistent behavior between the two SRS read lengths (Figure 2-B).

Finally, we compared the observed genera across all samples. The number of detected genera increased by 2.3-fold with NEB and by 4-fold with TruSeq when using 250PE reads compared to 150PE reads (Figure 2-C1), supporting the differential abundance findings between the two sequencing lengths. Additionally, the relative abundance profiles showed a greater proportion of genera classified as “other” with the 250PE sequencing, indicating an increase in taxonomic diversity, especially for the rare and low-abundance genera (Figure 2-C2). As expected, the PacBio dataset exhibited the lowest diversity due to its lower total read count (Figure 2-C1).

### Assessment of protein prediction and annotation

Principal coordinate analysis (PCoA) of beta diversity revealed a significant difference across NGS library preparation kits (Figure 3-A and B) for both 150PE (adonis2, R^2^ = 0.371, p = 0.001) and 250PE (adonis2, R^2^ = 0.322, p = 0.001) runs in the total detected proteins per replicate. No significant difference in beta diversity was detected in the NEB and Ovation datasets, regardless of the DNA input amount used for library preparation (data not shown).

**Figure 3.**
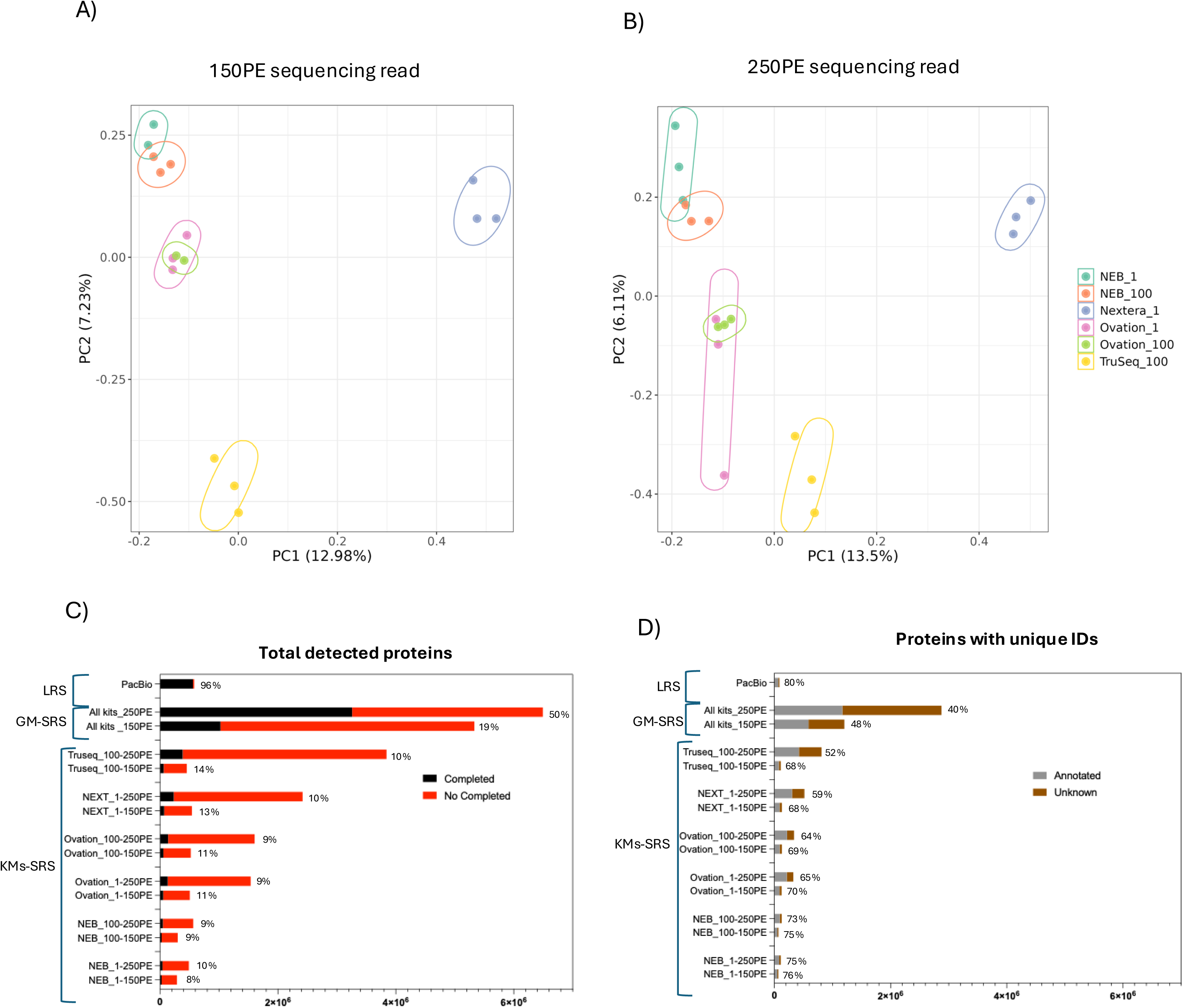
Comparison of library preparation kits and sequencing technologies for protein recovery. A and B) Principal Coordinate Analysis (PCoA) based on Bray-Curtis dissimilarity of detected gene profiles for each Illumina library preparation kit and DNA input replicate in the 150PE and 250PE runs. Each point represents a single sample and is colored by library preparation kit. Differences among groups were tested by PerMANOVA (R^2^ and p-value; 999 permutations). C) Bar plot showing the total number of detected genes (completed and fragmented) across the different co-assemblies (KMs and GM) and sequencing technologies (Illumina SRS at different read lengths and PacBio HiFi LRS). Fragmented proteins were exluded to reduce artifacts; only contigs >500 bp and proteins >100 bp (33 amino acids) were retained. Values indicate the percentage of complete proteins relative to the total number of detected proteins. D) Bar plot showing the number of unique proteins (annotated and unknown). Values indicate the percentage of annotated proteins relative to the total number of unique proteins.

For each KMs, in the 150PE mode, we observed only small differences in the total number of predicted proteins identified across the various kits, as shown in Figure 3-C. In contrast, the 250PE mode significantly enhanced protein recovery and gene completeness, a result consistent with the assembly metrics reported in Table 3. The TruSeq dataset, in particular, showed the most substantial improvement, with protein recovery increasing nearly tenfold. It also allowed the detection of approximately half of the non-completed proteins and around 10% of the completed ones when we compared against the GM catalogue (with six times higher sequencing depth). The rest of the kits also showed performance that reflected the assembly statistics listed in Table 5 and the contig distribution in Supplementary Figure 2, with the protein detection worsening with the decline in assembly and contig quality.

In the case of the LRS dataset, the total number of detected genes was in a range similar to those of the 150PE assemblies. However, the length of the assemblies allowed the retrieval of the complete sequence of 96% of the recovered genes.

When analysing the number of uniquely annotated proteins, in the 150PE run mode, we did not observe strong differences across the kits (Figure 4-D). In the GM, we observed a sharp increase in the total number of unique protein IDs, as expected due to the 6-fold increase in sequencing depth. However, the same libraries run at 250PE showed a significant enrichment in the total number of unique proteins detected, especially for the kits with better assembly parameters, such as TruSeq and Nextera, followed by the Ovations. Additionally, for the TruSeq libraries, the number of unknown proteins increased from 34,990 in the 150PE libraries to 389,567 in the 300PE libraries at the same sequencing depth, representing more than a tenfold increase. Finally, for the GM at 250PE the number of unknown proteins had the highest relative increase. In the case of the LRS results, the total number of unique proteins was in the same range as the 300 bp KM datasets, although the percentage of the unknown proteins was generally lower. This could be related to the features of the long-read sequencing technology, *per se,* that primarily catch the most abundant organisms.

**Figure 4.**
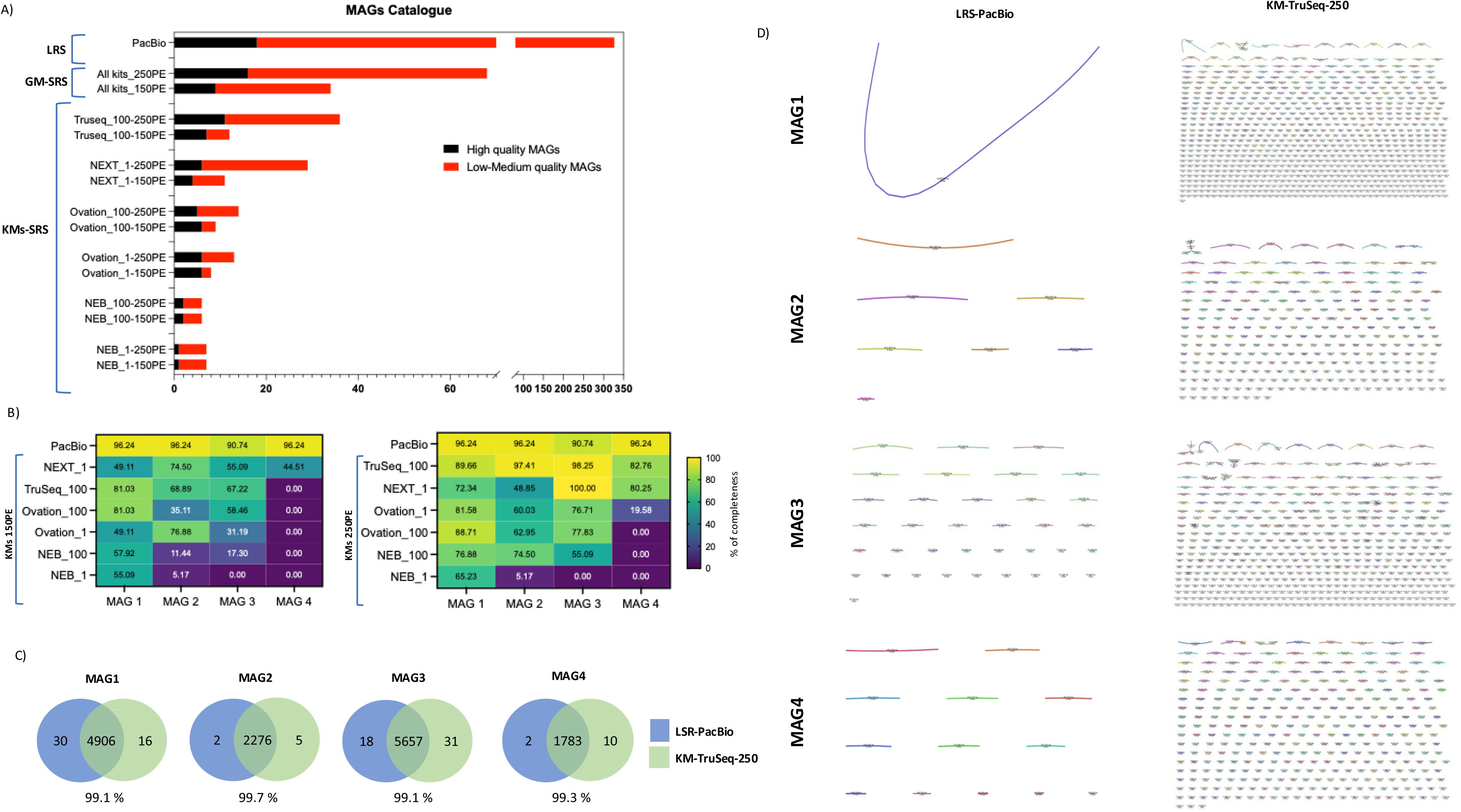
Comparison of library preparation kits and sequencing technologies for the MAG catalogue. A) Bar plot showing the total number of recovered MAGs (high and low-medium quality) across co-assemblies (KMs and GM) and sequencing technologies (Illumina SRS at different different read lengths and PacBio HiFi LRS). MAG quality thresholds were defined as follows: low quality, completeness <50% and contamination <10%; medium quality: completeness > 50% and contamination <10%; high-quality: completeness >90% and contamination < 5%. B) Heatmap showing MAG completeness (CheckM) for four new MAGs in KM Illumina libraries compared to PacBio. C) Comparison of orthologous protein clusters among sequencing strategies using OrthoVenn3. The diagram displays the number of unique and shared orthologous clusters identified in MAGs recovered from Illumina SRS KM-Truseq-250PE and PacBio HiFi LRS. Shared clusters represent conserved functions across datasets, while unique clusters may reflect method-specific functional recovery or assembly biases. OrthoVenn3 was run with default parameters (E-value cutoff 1e-5; inflation value 1.5). D) Assembly graph representations for the four new MAGs from Illumina SRS KM-Truseq-250PE and PacBio HiFi LRS technologies generated by Bandage.

### Metagenome-Assembled Genomes (MAGs)

We were also interested in measuring the effect of SRS versus LRS in the definition of MAGs. We expected LRS to generate higher-quality results due to the anticipated reduction in ambiguity during assembly. We determined that, despite the difference in sequencing depth, LRS could bin five times more MAGs than the GM assembly and almost ten times more than the best KM (TruSeq-250PE). When we compared the KM assemblies at 150PE, we observed the predicted trend, as in the protein catalogue, that the shortest library inserts produced the least MAGs (Figure 4-A). Although the total number of MAGs varied only modestly (from six MAGs in the NEB to 12 MAGs in the TruSeq at 150PE), the high-quality MAGs increased from 15% in the NEB kits to 58% in the TruSeq when measured at the same sequencing depth at 250PE. Across the KMs, the MAGs absolute recovery (low, medium, and high quality) was substantially higher in the TruSeq-250PE and Nextera-250PE. Notably, the LRS-derived MAG results were obtained using 40- and 13.7-fold less data than the GM and the KM-TruSeq-250PE, respectively (Table 5). Surprisingly, the number of high-quality MAGs was in the same order of magnitude among the three datasets, ranging from 18 in the PacBio to 16 in the GM and 11 in the KM-TruSeq-250PE (Figure 4-A).

### Characterization of novel MAGs matching speciation threshold

The combination of the results from CheckM and GTDB-Tk in the LRS dataset identified four high-quality MAGs that showed an average nucleotide identities (ANIs) value under 95%, which represents the standard speciation threshold (Jain et al. 2018). These results suggested these MAGs could be potential novel species. We then set out to verify the possibility that those were also present in the dataset generated by the different kits used for the SRS platform, possibly strengthening the identification of new species.

We ran a phylogenetic analysis of these new highest-quality bins from the LRS dataset to describe them taxonomically (Supplementary Figure 3). We identified four taxa that, based on their taxonomic novelty and lack of cultured representatives, could be classified as *microbial dark matter*. These included: (1) g JABFRZ01 (f GCA-002686595, Planctomycetota), (2) g VAYN01 (f 70-9, Actinobacteriota), (3) an unclassified genus within Pirellulaceae family (Planctomycetota), and (4) g UBA11591 (f Vicingaceae, Bacteroidota). All four MAGs showed classification only at high or intermediate taxonomic ranks (family or above), with placeholder genus names or no genus assignment at all, suggesting that they could represent phylogenetically novel bacterial lineages. Their detection highlights the presence of underexplored microbial diversity in the sample.

Then, we calculated the individual percentage of ANI and the MASH distance between the LRS and the respective KM bins (Supplementary Table 1) to verify the presence of similar bins in the SRS KMs. Once identified, we compared their percentage of completeness (Figure 4-B) against the LRS data. The KM-TruSeq-250PE combination achieved the highest completeness percentages and showed performance metrics most comparable to those obtained with the LRS approach (Supplementary Table 1 and Table 6). Overall, the long-read strategy produced better metrics and quality for these four MAGs, even though the total assembly data used (Table 5) was more than ten times larger in the KM-TruSeq-250PE. The number of contigs associated with each MAG was extremely different between the two technologies, most likely resulted from a higher level of fragmentation in the KM-TruSeq-250PE. As Figure 4-D and Table 6 show, the number of contigs ranged from one to 37 in LRS, whereas in the SRS KM-TruSeq-250PE, it spanned from 279 to 833. These results further reinforce the status of long-read sequencing as the method of choice for the recovery of high-quality MAGs. Despite the high fragmentation observed in the KM-TruSeq-250PE SRS bins, OrthoVenn analysis revealed a strong overlap in orthologous clusters between KM-TruSeq-250PE SRS and PacBio LRS MAGs, with similarity values ranging from 99.1% to 99.7% (Figure 4-C). Our functional analysis with AntiSMASH 8.0 identified 38 biosynthetic gene clusters in the PacBio LRS and 46 in the KM-Truseq-250PE SRS bins, including common types such as NRPS, Type I PKS, and terpenes. Several clusters had no close match in the MIBiG database, suggesting the potential for novel natural product biosynthesis (Supplementary Table 2).

**Table 6.**
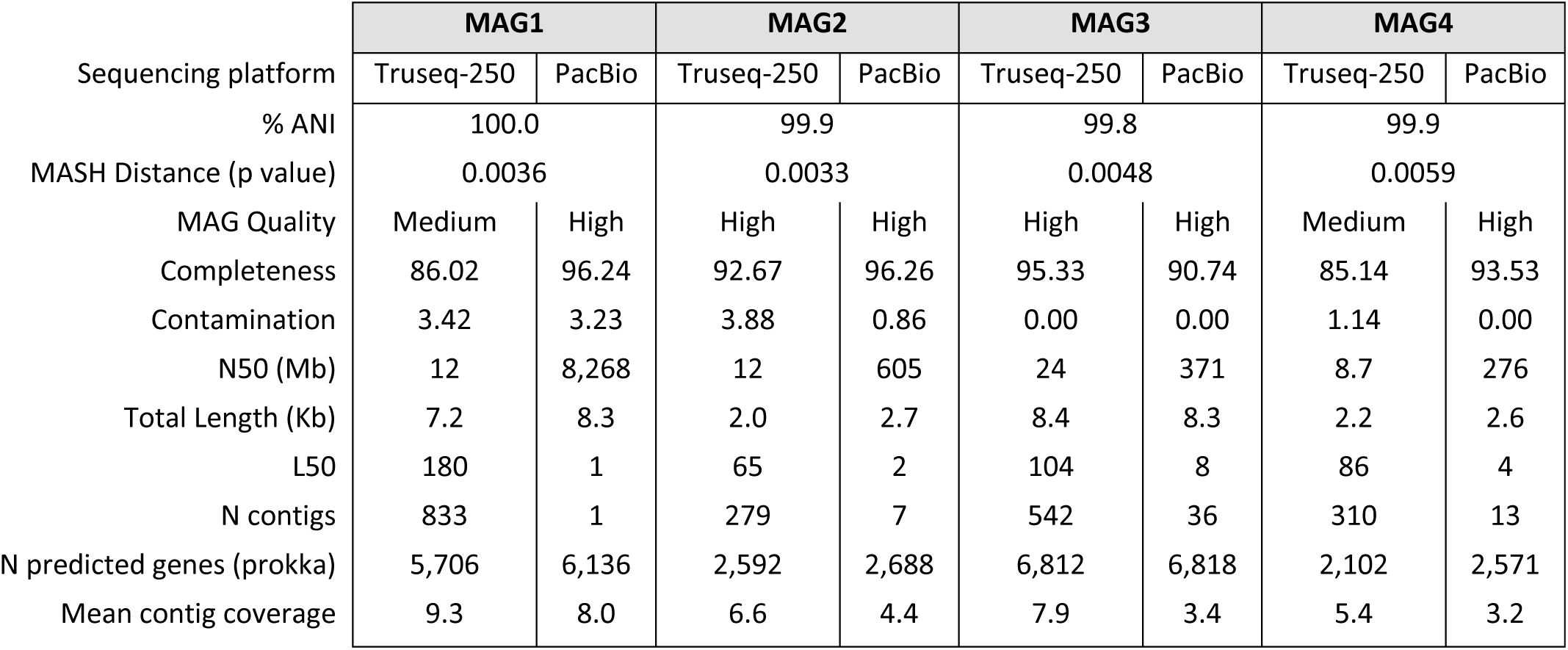
SRS KM-TruSeq-250PE and LRS PacBio MAG results comparison. % of ANI, p value of the MASH distance, MAG quality, % of completeness and contamination, N50 (Mb), total of the genome assembled (Kb), L50, number of contigs, number of predicted genes and mean contig coverage of the SRS KM-Truseq-250PE and LRS PacBio.

## DISCUSSION

The present study sought to maximize data acquisition and expand biological understanding in metagenomic analyses of highly diverse and complex samples. Our approach leveraged widely used library preparation kits and sequencing platforms, and we evaluated these methods with low DNA input, a common scenario for samples originating from extreme environments. Additionally, we utilised LRS to benchmark metagenome assemblies and bins as an internal reference, enabling us to evaluate short-read library preparation protocols directly on the community of interest rather than relying on low-diversity or artificial mock samples.

Overall, our comparison of different SRS kits revealed that the choice of library kit significantly influenced sequencing and assembly metrics, as well as taxonomic results, protein identification, and MAG recovery. In some of the cases, these effects were more evident when we coupled the kits with the 250PE sequencing read length. We found that the results were largely influenced by the fragment size distribution of the libraries generated by the kits, which affects the insert read length, that is, the fragment of the library that is effectively sequenced and carries biological information. This affected most of the subsequent steps, leading to adapter contamination, improved assembly metrics, lesser gene coverage and poorer MAG reconstruction. Furthermore, the tendency of SRS platforms to sequence preferentially shorter fragments reduced the amount of longer inserts present in the data (Ribarska et al. 2022).

In our opinion, the length of the effective insert size alone could not account for the differences observed between the kits. We confirmed this by bioinformatically constraining the downstream analysis to rely only on reads with lengths between 202 and 250 bp for NEB-100 replicates, thus matching the insert length of the TruSeq-250 kit, the best-performing library prep. Although this approach improved the assembly results compared to the original data, the computed metrics remained significantly lower than those of the TruSeq or Nextera kits at 250PE, at the same sequencing depth (Supplementary Table 3). Our analysis, though, does not exclude that by selecting longer fragments we biased the composition of those sequences, for example, excluding genes with a specific base composition less amenable to fragmentation.

Most genomic DNA-based sequencing library preparation protocols include a non-enzymatic shearing step. This method aims at breaking down the genome in an unbiased manner, thus creating a uniform distribution of DNA fragments independent of the base composition or other physical properties. Conversely, enzymatic fragmentation is biased since cut sites are never random and enzyme efficiency is susceptible to inhibitors. Consequently, physical shearing of DNA has remained the gold standard in sequencing. However, although it is efficient and consistent, it can be expensive and time-consuming. The method requires specialized instruments and consumables, adding significant cost and handling time (Head et al. 2014). Although solutions for the plate-based method already exist, those are not easily amenable to automation, thus making large-scale sequencing projects impractical for small labs and labor-intensive for larger ventures. Alternative methods available include endonucleases and transposase-mediated fragmentation and tagging (’tagmentation’). When the endonuclease-based fragmentation was introduced, concerns were raised that it might exhibit cut-site preference bias, therefore introducing artefacts (Knierim et al. 2011). However, advancements in commercial enzyme preparations addressed most of these issues using a cocktail of enzymes (Sato et al. 2019). Nonetheless, Tanaka et al. described that endonuclease-treated libraries exhibit a substantially larger number of recurrent artefactual SNVs/indels compared to those created through physical shearing (Tanaka et al. 2020). A previous report indicated that a fragmentase (developed by New England Biolabs) generates more artifactual indels than sonication or nebulisation. The number of indels produced by fragmentase was within a two-fold range of those generated by the physical methods (Knierim et al. 2011). In a similar study, Gregory et al. concluded that NGS library molecules produced by enzymatic fragmentation are not representative of the original DNA and are a byproduct of the enzymatic process. The authors found this observation consistent across different enzymatic fragmentation-based library preparation methods from other companies, suggesting that this is an inherent feature of the currently available commercial kits (Gregory et al. 2020). On the other hand, and consistent with observations related to enzyme-based fragmentation technologies, Gunasekera et al. reported biases in coverage, GC content, and de *novo* assembly quality metrics also when using tagmentation methods (Gunasekera et al. 2021). These findings may help explain the differences we observed in taxonomic assignment in the Nextera-prepared samples.

We recognise that library fragmentation could be optimized per each library kit and that in environmental samples, other aspects, such as the presence of humic acids and enzyme inhibitors, can affect the efficiency of any given kit, and each kit at a different level. However, ascertaining those effects goes beyond the scope of our study, requiring a sophisticated experimental design, material, time and funds.

In metagenomics analysis, *de novo* assembly is a critical step that will dictate the downstream genomics results, influencing the protein catalogue and quantity and quality of MAGs (Delgado and Andersson 2022; Kim et al. 2022). We used Megahit, which is a de Bruijn graph-based *de novo* assembler, that can lead to graphs that are complex to interpret since this method is affected by low or uneven coverage or sequencing errors, thus affecting the accuracy of contigs and scaffolds used for genome assembly (Ghurye et al. 2016). Sequencing bias at the library preparation level could be a significant problem for existing *de novo* assembly algorithms. De Bruijn-based assemblers such as Megahit use an average coverage cutoff threshold for contigs to prune out low coverage regions, which often contain more errors, thus reducing the complexity of the underlying de Bruijn graph. It also significantly affects the effective length and number of contigs in the final assembly, impacting the results (Li et al. 2015). We found that the percentage of high-quality MAGs is higher in the kits that use sonication as compared to the DNA enzymatic fragmentation step. This is in accord with Gregory and Tanaka’s observations of higher error rates on SNV/indels introduced by the enzymatic approach (Gregory et al. 2020; Tanaka et al. 2020). This is also likely to introduce more noise that affects the binning process, originating in higher contamination rates and fewer high-quality MAGs.

We speculate that the better results observed in the TruSeq kit could be a combination of different factors. The method requires a physical shearing step that, as we have discussed previously, provides a more uniform DNA fragment distribution, better suited for SRS (Head et al. 2014). The size selection step removes the shortest and longest fragments and improves the homogeneous clustering on the NGS platforms (Bronner and Quail 2019). In our study, the Ovation kits, whose protocol did not require a size selection step, performed worse than the TruSeq kits, suggesting that this process may positively impact the subsequent steps. The library prep protocols have many phases, largely divided in end-repair, A-tailing, ligation, indexes addition, and final amplification. At any of those, the difference in reagents among the manufacturers may have affected the quality of the final libraries. As an example, we found that TruSeq library kits exhibited lower duplication rates, even without reducing the number of PCR cycles.

Duarte et al., described that sequencing depth is crucial for gene discovery in marine microbial metagenomes. The authors highlighted that the low-abundance and unknown genes can be retrieved mainly using ultra-deep sequencing (Duarte et al. 2020), a factor that determines a significant cost increase. Our study suggested that the number of reads may not be the main parameter that impinges on the protein and MAG recovery in a complex environmental biome. We observed a significant improvement in the assembly metrics, even with the same number of reads, especially when the libraries had an optimal size distribution for sequencing at a 250PE length. When we employed the 250PE sequencing length, the assembly results in the TruSeq and Nextera were enhanced, potentially due to almost double the insert size that allowed the assembly algorithm to fill genome gaps, therefore delivering more accurate genomic information for the next steps in the analysis. This result suggests that increasing the sequencing length could be an excellent strategy for finding novel proteins and unknown taxa in complex environments.

Beyond NGS, as expected, third-generation LRS provides better assemblies, resulting in the recovery of high-quality MAGs and a higher percentage of fully assembled genes even using ten times less sequencing data (Meslier et al. 2022; Kim et al. 2022). However, the overall count of detected proteins was significantly lower, although assembling more data could certainly contribute to improving protein detection. It is currently impractical to compare the number of proteins detected by the KM-TruSeq-250PE to the one detected by PacBio since the depth of sequencing is largely in favor of the SRS method. This is likely to change in the near future, as continuous improvements in the sequencing output and cost reductions of long-read technologies become available. Despite this, the completeness of the proteins detected by the NGS-based kit was only 40%, whereas LRS reached 80%. It is well-known that PacBio sequencing technology struggles with gene recovery and the assembly of organisms present in a low-abundance (Kim et al. 2024). This limitation can impact studies of highly fragmented and complex biomes such as soil (Alteio et al. 2020) or studies aimed at characterizing the rare biosphere (Jin et al. 2022). Although using third-generation LRS for metagenomic experiments offers substantial advantages, particularly for genome assembly and MAG recovery, it is also important to consider the cost and the stringent sample requirements. These aspects become particularly critical for large-scale projects, especially in extreme biomes that include samples with scarce DNA and genome integrity. An interesting approach is to include long-read sequencing (LRS) for a subset of samples within a project to generate an internal reference MAG catalogue, enabling robust benchmarking of short-read (SRS) assemblies on real environmental data (Sanders et al. 2019). This strategy could provide a realistic framework for evaluating sequencing and assembly performance in complex and novel microbial communities, and for guiding future efforts aimed at maximizing overall MAG recovery.

Our key discovery indicates that the TruSeq Illumina library preparation kit, when combined with a 250PE sequencing read length, can yield results that closely align with those obtained using long-read technologies for high-quality MAGs in a highly diverse environmental sample. Despite the fragmentation typical of short-read assemblies, the conservation of orthologous clusters found in the MAGs with long-read assemblies indicates reliable recovery of key functional genes and pathways. Additionally, this combination (TruSeq and 250PE) provides the highest number of protein recoveries among the SRS kits, offering the most holistic view of community diversity and function, an outcome not achievable with other library preparation or sequencing strategies. In our opinion, the Nextera kit is also a suitable option, particularly for low-input samples, showing competitive performance in both protein and MAG recovery. However, the taxonomic discrepancies observed in Nextera-prepared samples could raise concerns that warrant further investigation. Additionally, the streamlined nature of the Nextera workflow makes it well-suited for automation. Nevertheless, it is important to note that tagmentation-based technologies can be susceptible to the presence of inhibitors, particularly present in environmental samples, which may negatively impact the library preparation process.

## CONCLUSION

To our knowledge, this study provides the first direct assessment of 2×150 bp versus 2×250 bp Illumina short-read metagenomic sequencing across multiple library preparation kits, using the same complex environmental sample on the NovaSeq 6000. The findings highlight that read length can substantially influence the resolution of metagenomic assemblies and the reconstruction of microbial genomes, providing a valuable reference for future large-scale metagenomic projects. Our study concludes that selecting the appropriate library preparation method, sequencing length and technology is crucial for enhancing outcomes of metagenomic studies and deciphering unknown microbial information. The optimal approach, however, depends on factors like sample type, DNA quantity, budget, and crucially, your study’s specific aims. Our study highlights the critical importance of rigorous laboratory practices, thorough library quality control, and the careful selection of appropriate methodologies to enhance the detection of proteins and MAGs. These factors are essential for ensuring data comparability across samples and studies, particularly in large consortium projects and meta-analyses. This is especially relevant for datasets generated by commercial sequencing facilities, where details of library preparation methods are often insufficiently reported by the authors (Li et al. 2017; 2020).

Overall, our findings suggest that some current library preparation and sequencing technologies still limit our ability to fully capture microbial information in certain environmental samples, particularly those from complex, unknown and underexplored environments. This indicates that a vast amount of untapped genetic content remains hidden, paving the way for future discoveries. Importantly, re-sequencing existing samples using a longer sequencing read length becomes a viable option, saving setup costs and allowing continued deciphering of the dark matter in complex biomes without needing new and costly sampling experiments.

## Acknowledgments

This work was funded by the Bioscience Core Lab operational fund.

## SUPPLEMENTARY MATERIAL

**Supplementary Figure 1.** Library preparation results. A) Bioanalyzer profiles showing the size distribution for each Illumina library preparation kit. B) Final library concentration and number of PCR cycles used for each kit, following the manufacturer’s protocol. Library concentration was measured from 1 µL of the final library using the Qubit HS dsDNA assay. Bold numbers indicate the enrichment PCR cycles used for each Illumina SRS kit. C) Average library fragment size for each kit, calculated using the Bioanalyzer analysis software.

**Supplementary Figure 2.** Contig length distribution for the Illumina KM co-assemblies (A, 150PE; B, 250PE) and the PacBio assembly (C). Histograms of contig lengths were generated to assess and visualize assembly quality. The x-axis shows contig length (bp), and the y-axis shows the frequency (total number of contigs) within each length bin.

**Supplementary Figure 3.** Closest taxonomy assignment and phylogenetic placement for the four new MAGs from the PacBio dataset. For each genome, an appropriate outgroup was selected for phylogenetic analysis: PacBio_MAG1, p_Planctomycetota; PacBio_MAG2, g_Rubrobacter; PacBio_MAG3, f_Planctomycetaceae; PacBio_MAG4, o_Bacteroidales.

**Supplementary Table 1.** Comparison of MAG metrics between Illumina SRS KM-TruSeq-250PE and PacBio HiFi LRS assemblies. Reported values include ANI (%), MASH distance (p value), MAG quality, completeness (%) and contamination (%), N50 (Mb), total assembled genome size (kb), L50, number of contigs, number of predicted genes, and mean contig coverage.

**Supplementary Table 2.** Comparative analysis of biosynthetic gene clusters (BGCs) predicted by antiSMASH 6 in MAGs recovered from Illumina SRS KM-Truseq-250PE and PacBio HiFi LRS bins. The table summarizes the number and classes of BGCs identified (e.g., NRPS, PKS, RiPP-like, terpene, and other specialized metabolites). Predicted clusters are annotated based on similarity to known BGCs in the MIBiG database; percent similarity indicates the degree of homology.

